# Differential effects of aging and Alzheimer’s disease on microemboli clearance in a mouse model of microinfarction

**DOI:** 10.1101/2025.09.30.679680

**Authors:** Fiona O. Haugen, Gergely Silasi

## Abstract

**Background:** Cerebral microinfarcts often occur as a result of microvessel occlusion and are prevalent among dementia patients and the aging population. Detailed studies on the timecourse of microvascular occlusions indicate that endogenous mechanisms exist to re-canalize occluded vessels. One recently discovered mechanism is angiophagy, where vessels engulf and expel microemboli, thus mitigating damage caused by micro-occlusions. While several previous studies have shown that angiophagy occurs in rodent models, the frequency and timing of this process is not well characterized. In addition, there is limited data on the impact of aging on angiophagy, or the occurrence of this process in clinically relevant diseases such as Alzheimer’s disease.

**Methods:** To further study the timecourse of angiophagy, we induced micro-occlusions in young, aged and 3xTg Alzheimer’s mice via injection of 20μm microspheres into the carotid artery. Mice were sacrificed on day 3, 7 or 14 and the brains were processed for brain-wide localization of microspheres and quantification of angiophagy.

**Results:** We found the largest number of microspheres in the neocortex, yet when accounting for region size, microspheres were more evenly distributed across brain regions. When quantifying angiophagy in young non-diseased mice, we found that approximately 43% of microspheres had extravasated from the vessel by day 14. This process was delayed in aged mice, with only 10% of microspheres extravasated by day 14. Moreover, in young 3xTg Alzheimer’s mice, we found the rate of angiophagy to be more efficient at day 14 compared to non-transgenic controls, with 47% and 43% of microspheres extravasated, respectively. A similar trend was observed in aged Alzheimer’s mice, in which 38% of microspheres were extravasated by day 14 in 3xTg mice, compared to only 30% in non-transgenic controls.

**Conclusions:** Taken together, we find that while aging impairs the process of angiophagy, Alzheimer’s mice exhibit a paradoxical increase in the rate of microsphere extravasation.

**Graphical abstract:** 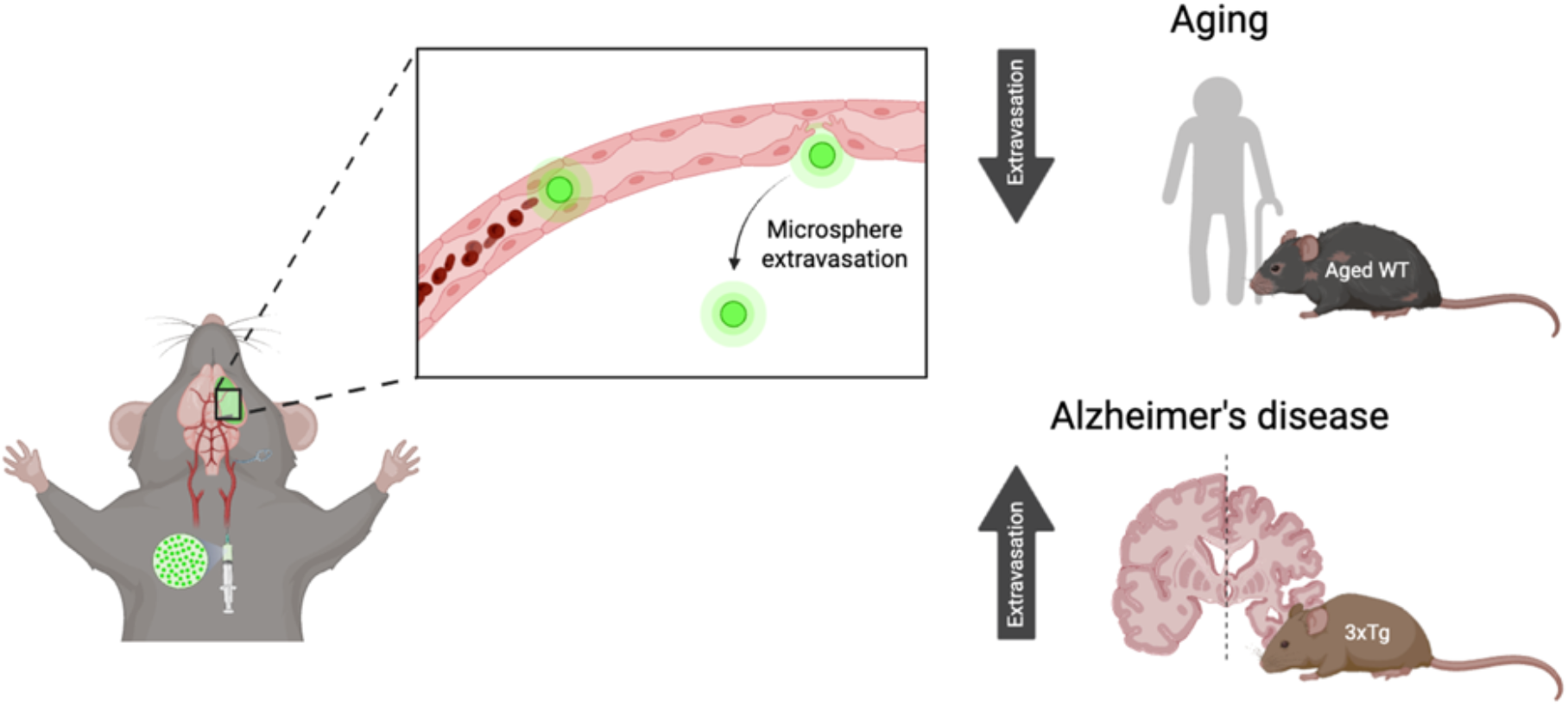

## Introduction

Occlusion of cerebral microvessels results in the formation of microinfarcts (MIs), which range from 100*μ*m to 3mm in size^1^. It is hypothesized that MIs accumulate overtime, with individuals amassing hundreds to thousands over a lifetime^1^. Although seemingly insignificant, MIs can contribute to cognitive decline and are most common among the aging population^1^. MIs are found in approximately 62% of patients with vascular dementia, 43% of patients with Alzheimer’s disease (AD), and in 24% of individuals 75 years or older^2^. Notably, AD patients with microinfarcts have more severe cognitive impairments than AD patients without microinfarcts^3^.

Despite the high incidence of MIs in subpopulations of individuals, the majority of intravascular emboli are thought to be in fact cleared out by hemodynamic forces or chemical breakdown via thrombolysis^4^. Although these clearance mechanisms hold great therapeutic promise, they are not always effective. For example, in Alzheimer’s disease, areas of hypoperfused brain regions lack sufficient blood flow for hemodynamic forces to act effectively^5,6^. Interestingly, recent experiments have suggested a third method of emboli clearance known as angiophagy^7^. This mechanism relies on a form of microvessel remodeling where emboli are engulfed and translocated out of the vessel into the surrounding parenchyma, thus facilitating the re-establishment of blood flow^7^. Although the exact structural and molecular mechanisms of angiophagy are still unknown, Lam et al. noted that endothelial projections engulf occluding emboli for translocation, and this process may be enabled by heightened levels of matrix metalloproteinases (MMP) that are detected at sites of extravasation^4,7^. Initial morphological features of angiophagy (lamellipodia projections) have been shown to initiate as early as 3 hours after vessel occlusion^4^. While several additional studies have investigated the timecourse of angiophagy in young murine micro-occlusion models, the results have been variable^7-9^. Moreover, one study reported that angiophagy was decreased by ∼35% in aged mice, however this effect has not been further investigated^7^.

Our current study compared the brain-wide localization of microspheres in young and aged non-transgenic and Alzheimer’s 3xTg mice. To address the discrepancies in the rate of reported angiophagy in previous studies, we quantified angiophagy across multiple brain regions in both young and aged mice up to 14 days after microsphere injections. Our study confirms that angiophagy is impaired in aging by showing that the rate of extravasation in aged mice was significantly delayed. In addition, our experiments showed that in Alzheimer’s 3xTg mice there was an increase in microsphere angiophagy, despite clinical reports of a higher incidence of microinfarcts in individuals with Alzheimer’s disease^2^.

## Methods

### Animals

All experimental protocols were performed in accordance with the University of Ottawa’s Animal Care Committee regulations. Male and female wildtype mice (n=54) with a C57BL6 background (RRID:IMSR_JAX:031823 and IMSR_JAX:007612) and Alzheimer’s disease mice (n=33) with strain-matched controls (n=37) with a B6;129 background (RRID:MMRRC_034830-JAX and IMSR_JAX:101045) were used. Mice were between 2 and 6 months of age for the young cohort and between 9 and 20 months for the aged cohort. Of the 127 total mice, 16 were excluded as a result of post-surgery mortality and 31 were excluded from total microsphere and/or angiophagy quantification as a result of unsuccessful surgeries (<50 microspheres) or poor vascular labelling.

### Experimental procedures

Surgical procedures were performed under isoflurane inhalation anesthesia mixed with oxygen (3.5% isoflurane-1L/min oxygen for induction, 2% for maintenance). Body temperature was maintained at 37°C with a heating pad. Microvessel occlusions were induced via the injection of 15um (diameter) polystyrene microspheres into the left carotid artery while the external carotid artery is clamped. Mice were euthanized on day 3, 7 or 14 post-surgery via injection of sodium pentobarbital (65mg/mL, 10mL/kg). For visualization of vessels, lectin and/or Evans Blue was injected intracardially and allowed to circulate for 5 minutes prior to brain extraction. Brain samples were post-fixed in 4% paraformaldehyde for a minimum of 24 hours prior to serial sectioning on a vibratome at 100*μ*m.

### Sample analysis

A semi-automated image analysis pipeline was used to quantify the number of microspheres brain-wide. Coronal brain sections (100*μ*m) were imaged under a fluorescent microscope (10x objective) with a motorized stage and images were stitched to re-create sections. To automatically identify microspheres in histological sections, the image classification software Ilastik was trained using manually annotated images. QuickNII was used to align the sections with the Allen mouse brain atlas, and finally Nutil was used to overlay the Ilastik segmentation files onto the atlas allowing for quantification and localization of the microspheres.

Angiophagy quantification was performed live under 20x magnification on a Ziess Axio Imager M2 fluorescence microscope. Microspheres were classified into one of three categories based on location relative to vessel: obstructing (in the vessel), going out of the vessel, and extravasated from the vessel. The anatomical brain region of the microsphere was also determined based on the overlapping atlas location for each microsphere.

## Results

### Microspheres primarily localize to the neocortex with the greatest density of microspheres in the thalamus

Using the semi-automated QUINT image analysis pipeline (Figure 2 A,B), we determined that although the total number of microspheres varied among individual subjects, the mean number of microspheres is generally comparable across experimental groups. The young NTG mice had the greatest number of microspheres (890.7±90.75 microspheres) and the aged non-diseased had the least number of microspheres (490.5±68.6 microspheres; Figure 2C).

**Figure 1.**
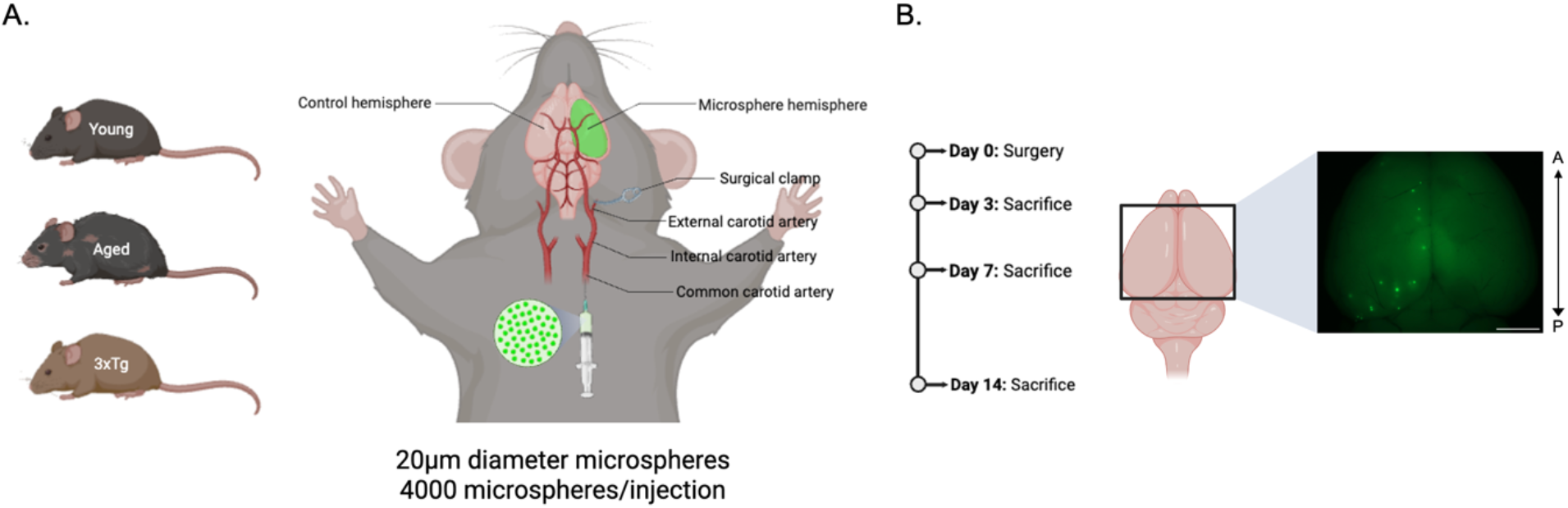
Experimental design. Mice are injected with 20μm (diameter) microspheres via the common carotid artery **(A)** and are sacrificed on day 3, day 7 or day 14 **(B)** following an intracardial injection of Evans Blue and/or lectin for vascular labelling. Scale bar 5mm.

**Figure 2.**
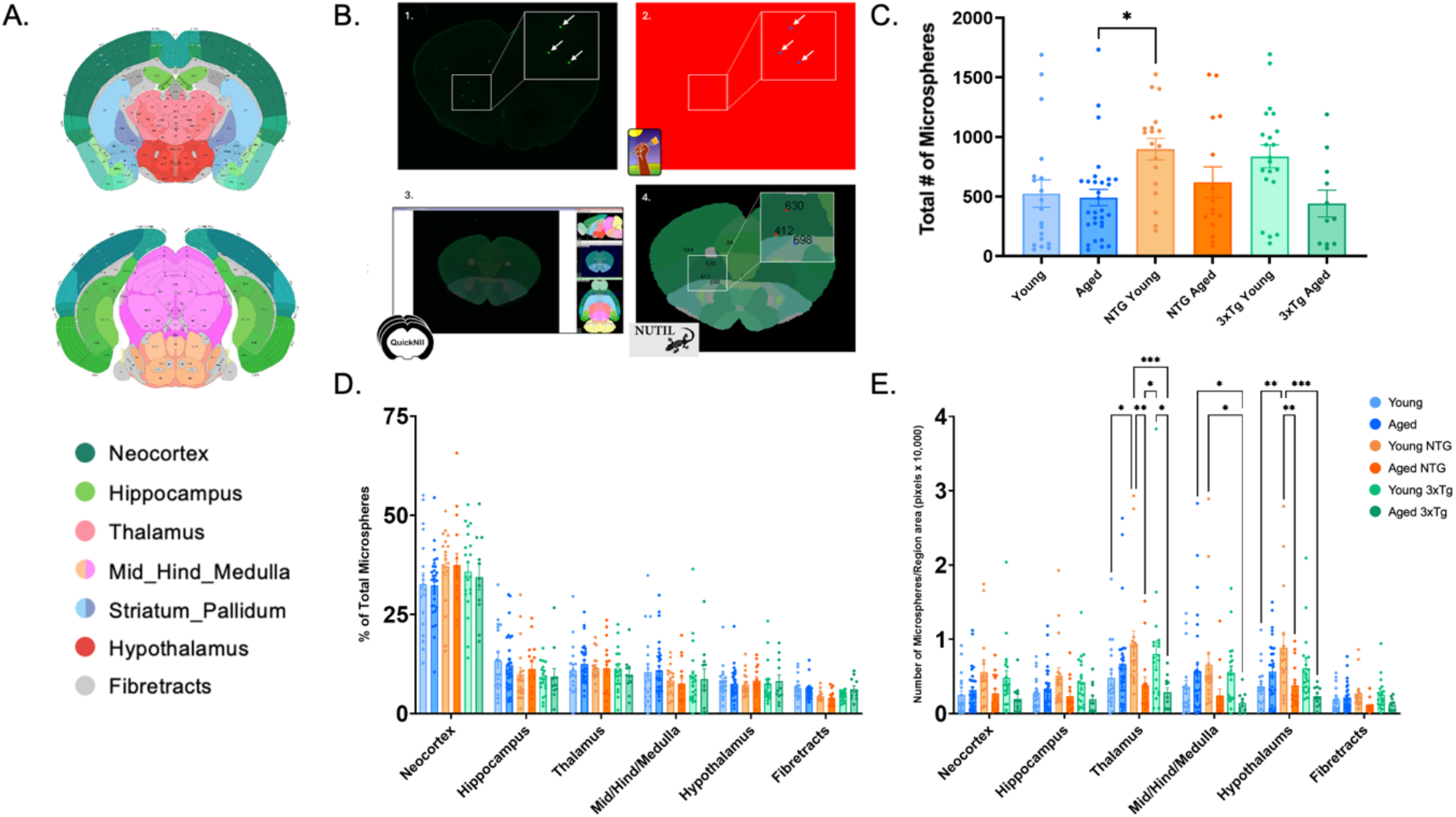
Microsphere localization for all experimental cohorts. Using a semi-automated image analysis pipeline, microsphere localization was determined in young non-diseased, aged non-diseased, young NTG, aged NTG, young 3xTg, and aged 3xTg mice. **(A)** Allen Mouse Brain Atlas for region legend. **(B)** QUINT image analysis pipeline. **(C)** Total number of microspheres across cohorts. **(D)** Regional microsphere localization. **(E)** Normalized microsphere localization. Regions not shown: olfactory, ventricular system, cerebellum. Data expressed as mean ± SEM. Two-way ANOVA with Tukey’s multiple comparison. ^***^p<0.001, ^**^p<0.01, ^*^p<0.05.

Aligning Nutil segmentation files of the coronal brain sections with the Allen Mouse Brian atlas allowed us to determine the regional distribution of microspheres. The greatest percent of microspheres localized to the neocortex, which was found to be consistent across all experimental groups (Figures 2D). When comparing the percent of microspheres within a brain region across each experimental group, there was no significant difference among groups (Figure 2D). After normalizing the number of microspheres by the area of each brain region, we found significant differences among groups in microsphere density in some structures (thalamus, midbrain, hypothalamus), however the only consistent effect was that the thalamus had the greatest density of microspheres across all experimental groups (Figure 2E).

### Microemboli extravasation is delayed in aged non-diseased mice

To determine the time course of angiophagy in our microsphere injection model, we quantified the process at three timepoints: day 3, 7, and 14. Microspheres were categorized into one of three groups depending on their location relative to labelled vessels: obstructing the vessel, in the process of going out of the vessel, and extravasated outside of the vessel (Figure 3A).

**Figure 3.**
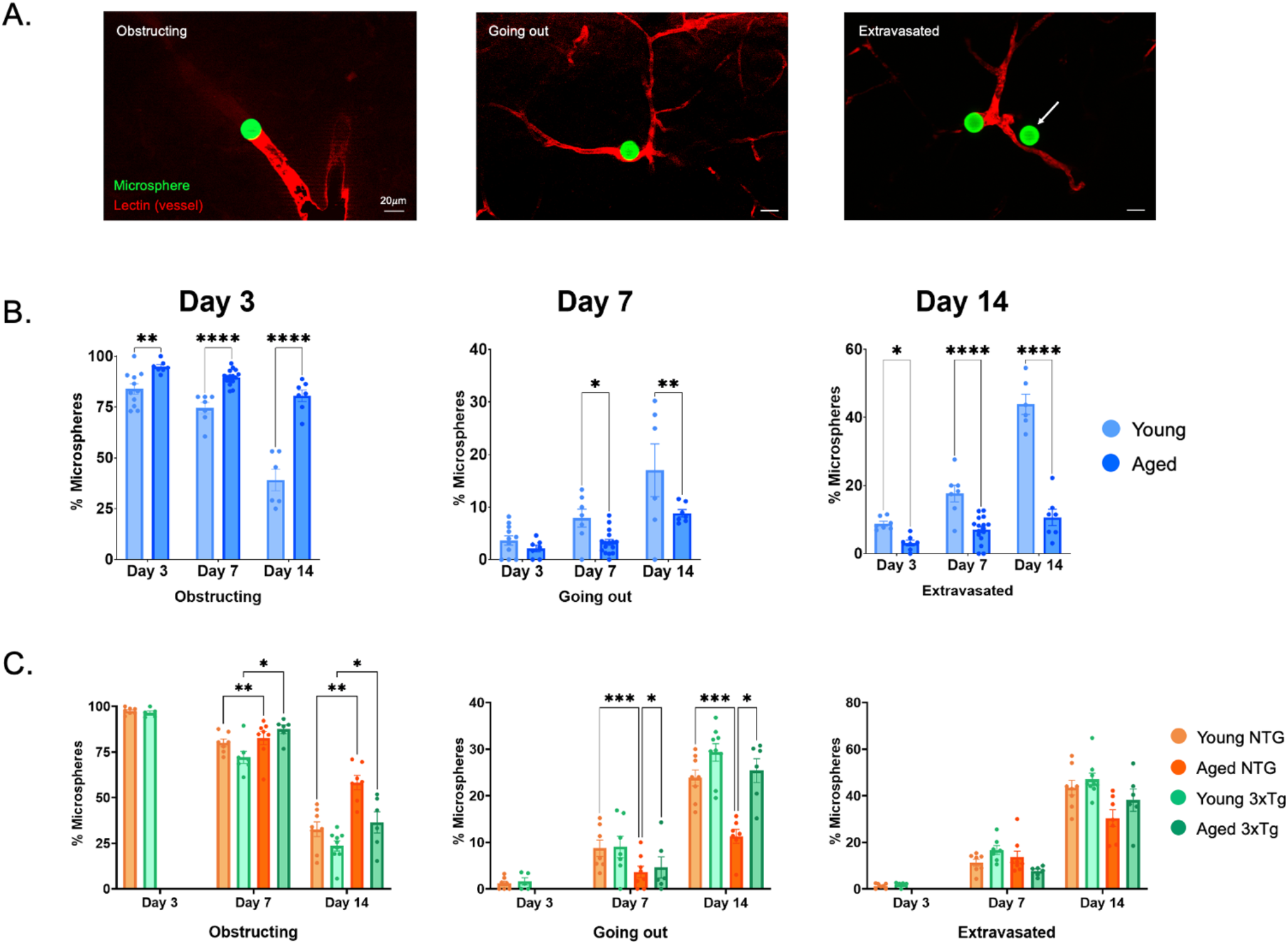
Quantification of angiophagy. **(A)** Representative images showing angiophagy categories used for quantification: obstructing, going out and extravasated. **(B)** Angiophagy quantification comparing young and aged wildtype mice. **(C)** Angiophagy quantification comparing young and aged NTG and 3xTg mice. Data expressed as mean ± SEM. Two-way ANOVA with Tukey’s multiple comparison. ^****^p<0.0001, ^***^p<0.001, ^**^p<0.01, ^*^p<0.05.

In the young NTG cohort, 43.85±2.95% of microspheres had extravasated by day 14 (Figure 3B). Across all time points, extravasation was significantly delayed in aged wildtype mice compared to young counterparts, with only 10.6±2.4% of microspheres extravasated by day 14 (Figure 3B). Angiophagy was also assessed within brain regions by sampling 5-20 microspheres per structure (neocortex, striatum, thalamus, hippocampus and white matter). While young animals showed a significant increase in angiophagy at day 14 across all brain regions examined, the aged group did not show a similar effect (Supplemental Figure 1).

### Alzheimer’s disease mice show increased angiophagy efficiency compared to non-transgenic controls

In order to assess the effects of Alzheimer’s pathology on the process of angiophagy, we performed the same surgery on young (prior to onset of amyloid and tau pathology) and aged 3xTg mice with age-matched non-transgenic controls. Comparable to our non-diseased cohort, we observed a similar trend of delayed angiophagy in aged 3xTg mice, as at both day 7 and 14 aged mice have a greater percentage of microspheres obstructing vessels compared to young 3xTg mice (Figure 3C).

Notably, we observed a general trend for 3xTg mice to be more efficient at the process of angiophagy, particularly in the aged group, at the day 14 timepoint. At day 7 and 14, the percent of microspheres going out of vessels was significantly greater in 3xTg aged mice compared to NTG age-matched controls (Figure 3C).

### Minimal correlations between microsphere load and the occurrence of angiophagy

Correlation plots were created to investigate a potential relationship between microsphere load and the rate of angiophagy. At day 3 and 14 in young mice and day 3 and 7 in aged mice, we observed a positive correlation, whereby a greater microsphere load correlates to a greater percent of extravasated microspheres, or increased occurrence of angiophagy (Figure 4A, C, D, E). Young mice at day 3 and aged mice at day 14 appear to have a general negative correlation, as a greater microsphere load correlates to less extravasated microspheres, or a decreased occurrence of angiophagy (Figure 4B, F). Notably, correlation between microsphere load and percent of extravasated microspheres is only significant in the aged non-diseased mice at day 3 with an r value of 0.8548 (Figure 4D).

**Figure 4.**
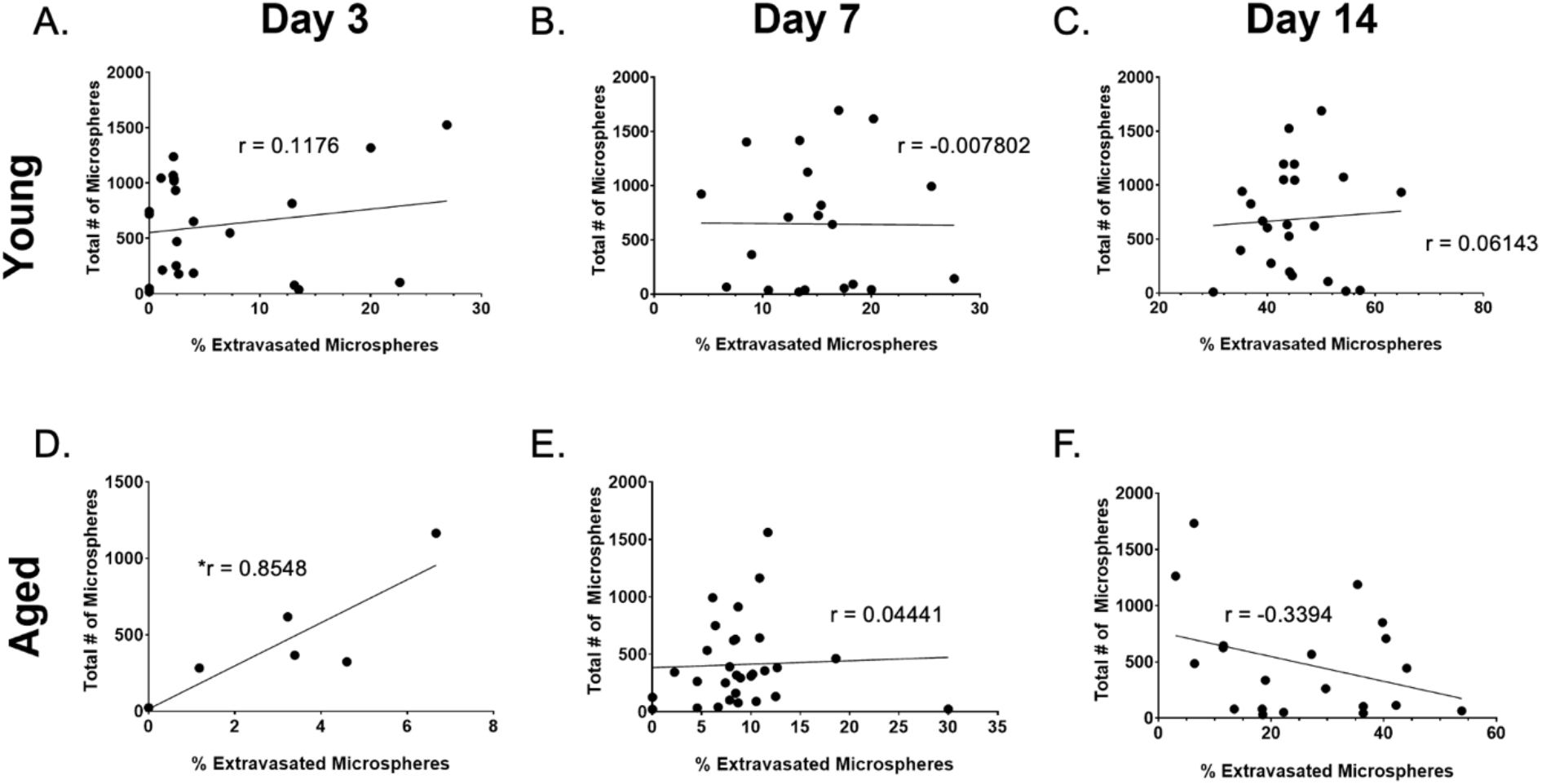
Correlation between microsphere load and extravasated microspheres. Correlation between microsphere load and extravasated microspheres in young mice **(A, B and C)** and aged mice **(D, E, and F)** at day 3, 7 and 14. ^*^ = significant (alpha = 0.05).

## Discussion

The diffuse microemboli model used in the current study is commonly employed to study microinfarcts and emboli clearance mechanisms^4,7-9^. Using the semi-automated QUINT analysis pipeline^10^, we were able to characterize this model by performing brain-wide quantification and localization of all microspheres in individual mice. Although the mean number of recovered microspheres varied across groups, we did not observe a significant effect of age or genotype, indicating that the initial micro-occlusion load is similar across groups.

Our previous work on this model has developed and characterized methods for quantifying the distribution of microspheres across brain regions^11,12^. Our current results confirm that the neocortex accumulates more microspheres than any other region, followed by the thalamus and hippocampus^11-16^. When the size of the brain region was taken into account, we found that the thalamus had a greater number of microspheres per volume of brain region for all cohorts, which is in line with our previous study showing that the thalamus had the highest density of microspheres^12^.

Microvessel occlusions are a common cause of microinfarcts, yet the brain has protective mechanisms in place, such as angiophagy, to clear microemboli and facilitate reperfusion^4^. Previous studies have presented angiophagy rates ranging from 18-80%, which may be explained by small sample sizes (40-195 microspheres quantified total for each timepoint) and the use of varying sizes and composition of microemboli^7-9^. Our goal was to further investigate the timecourse of angiophagy with a larger sample size, categorizing 10-100 microspheres per mouse, with 5-16 mice used at each of the three timepoints (mean of 8 mice quantified per timepoint).

Our results show that approximately 17% of microspheres are extravasated from vessels at the day 7 timepoint in young, non-diseased mice. This is in contrast to previous studies, which reported 78% or 58% of fibrin and cholesterol clots, respectively, or 33% of polystyrene microspheres extravasated at day 7^7,9^. One study even suggested 76% of microspheres had undergone angiophagy by day 7^8^. Our finding of approximately 43% microspheres extravasated at day 14 is comparable to the first study on angiophagy published by Lam et al. citing that just over 45% of microspheres (10-15*μ*m in diameter) had undergone extravasation at this time point^6^. Importantly, our results confirm that this relatively recently described phenomenon can be visualized and quantified following fluorescent polystyrene microsphere injections in mice.

Our histological processing pipeline also allowed us to localize each occlusion site to the Allen Mouse Brain common coordinate framework, thus allowing us to examine if the rate of angiophagy varied across brain regions. One possibility may be that angiophagy is initiated by ischemic signals or cell damage that is produced at occlusion sites, therefore regions with higher microsphere load, such as the neocortex, may have higher levels of angiophagy^17^. However, we found no significant difference in angiophagy across brain regions, suggesting that signaling mechanisms initiating this process are regulated globally across the brain, likely independent of the number of micro-occlusions or the degree of ischemia.

Previously, Lam et al. studied the effects of aging on the process of angiophagy and found that clot extravasation (fibrin or cholesterol) in aged mice decreased by just over 20% at day 4 and microsphere extravasation (polystyrene) decreased markedly by just over 30% at day 14 relative to young mice^7^. We present similar findings in aged mice whereby at day 14, microsphere extravasation decreased by just over 30%. In humans, it has been found that microinfarcts are present in 24% of non-demented individuals over the age of 75^2^. It may be the case that vascular alterations seen in aging play a role in the observed decrease in the rate of angiophagy, which may subsequently lead to the development of MIs. For example, it is possible an increase in basal lamina thickness and collagen deposition, or arterial stiffening caused by a decrease in elastin expression make it more difficult for endothelial remodeling to occur and subsequently the engulfment and translocation of microemboli^18-20^.

Moreover, microinfarcts are commonly observed in those with Alzheimer’s disease, presenting in 43% of AD patients^2^. We hypothesize that vascular changes associated with AD, such as arterial stiffening due to cerebral amyloid angiopathy (CAA), may impair this process of angiophagy^21^. To test our hypothesis, we used a 3xTg mouse model of AD, which have been found to have pathological differences beyond 11 months including decreased arterial flow, decreased density of penetrating arterioles, and importantly for studying angiophagy, vascular amyloid *β* accumulation and thickened basement membrane, which may impact endothelial remodeling^22-24^.

Based on previous literature, severe AD pathology and vascular differences are not predicted to be present in the young 3xTg cohort (2-6 months)^22-25^. Our results are in agreement with this as we observed comparable levels of angiophagy between young NTG and young 3xTg mice. Although not significant, there was a general trend at day 14 for 3xTg mice to have fewer obstructing microspheres and more microspheres in the going out and extravasated stages compared to young NTG mice. A possible explanation for this is the hypothesized role of matrix metalloproteinases (MMPs) in the process of angiophagy. Gelatinases A (MMP2) and B (MMP9) are commonly studied in the brain and play a role in the disruption of blood brain barrier, whereby MMP-mediated blood brain barrier (BBB) disruption is a result of MMPs attacking the extracellular matrix, basal lamina and tight junctions of endothelial cells^26^. In studying angiophagy, Lam et al. found increased gelatinolytic activity at sites of occluded vessels compared to unoccluded vessels and a decrease in angiophagy with the addition of a MMP2/9 inhibitor^7^. This points to the possibility that MMPs play a role in angiophagy. Importantly, MMPs have also been implicated in AD and may facilitate the process of amyloid β clearance^27,28^. It has been found that MMPs are endogenously induced by amyloid molecules in blood vessels, astrocytes, and microglia^28^. Moreover, APP/PSEN1 transgenic mice have shown to have increased MMP2 and MMP9 concentrations in astrocytes around amyloid plaques^28^. Since the presence of amyloid β has been shown to induce expression of MMP2/9, it is possible that 3xTg mice display heightened levels of MMP2/9 activity. This could subsequently account for the increase in the percent of microspheres going out and extravasated in young 3xTg mice compared to young NTG mice.

We initially hypothesized that the progression of Alzheimer’s disease pathology in our aged AD mice would impair the process of angiophagy as a result of CAA or thickened basement membrane^22^. Interestingly, we find that aged 3xTg mice were more efficient at angiophagy as at day 7 and 14, significantly more microspheres were in the going out stage compared to aged NTG mice. Higher levels of MMPs have been associated with late-stage Alzheimer’s disease, so it may be the case that 3xTg are more efficient at the process of angiophagy as a result of greater MMP expression at this age^29^. Moreover, it may be the case that clearance mechanisms (washout and angiophagy) work in concert, and when one mechanism is impaired, the other upregulates to compensate. Emboli washout via blood flow may be impaired in AD as a result of observed hypoperfusion, and in turn we see a compensatory increase in angiophagy in our AD mouse model^5^.

Lastly, we wanted to explore the possibility that the number of microspheres present in the brain impacted the rate of angiophagy, which could explain previously recorded discrepancies in the timeline of angiophagy. We created correlation plots of the percent of extravasated microspheres and the number of microspheres in each mouse across all cohorts and time points. The aged non-diseased cohort at day 3 was the only correlation found to have a significant relationship as a higher number of microspheres were linked with a higher percent of extravasation, however our sample size was relatively small for assessing this relationship.

In conclusion, we characterized microsphere distribution using an open-source, semi-automated image analysis pipeline. Based on our data, we offer a comprehensive timeline of angiophagy in aged and Alzheimer’s disease mice, while elaborating on the current angiophagy timelines in young, non-diseased rodents in an effort to reconcile discrepancies in previous literature. We found that angiophagy is delayed in aged mice and interestingly, AD appears to enhance the efficiency of the extravasation process. Furthermore, we are the first to compare angiophagy in different brain regions, as well as investigate a potential correlation between microsphere load and the occurrence of angiophagy. Defining a timecourse for angiophagy is important in understanding this process as a potential protective mechanism following microvascular occlusion. Moreover, future therapeutic modulation of angiophagy will require a detailed understanding of the timecourse of this process in the context of microinfarction as well as comorbidities such as Alzheimer’s disease.

## Supplemental

### Supplemental methods

#### Animals

| Cohort | Strain | Sex | Excluded | Died | Total Included |
| --- | --- | --- | --- | --- | --- |
| Young | Tmem119 | F = 15 | n = 14 | n = 5 | 24 |
|  |  | M = 28 |  |  |  |
|  | NTG | F = 10 | n = 1 | n = 0 | 22 |
|  |  | M = 13 |  |  |  |
|  | 3xTg | F = 13 | n = 8 | n = 0 | 21 |
|  |  | M = 16 |  |  |  |
| Aged | Tmem119 | F = 13 | n = 2 | n = 9 | 30 |
|  |  | M = 0 |  |  |  |
|  | ChR2 | F = 22 |  |  |  |
|  |  | M = 7 |  |  |  |
|  | NTG | F = 5 | n = 1 | n = 0 | 15 |
|  |  | M = 16 |  |  |  |
|  | 3xTg | F = 4 | n = 6 | n = 2 | 12 |
|  |  | M = 16 |  |  |  |

#### Microvessel occlusion model

##### Solution preparation

A fluorescent microsphere model of microinfarction was used, first described by Silasi et al^11^. A working solution of 3000-4000 microspheres/100*μ*L injection was prepared prior to surgery using the stock solution of fluorescent microspheres (FluoroSpheres polystyrene, Polysciences, Cat# 19096-2; 20um particles, 2.5% aqueous suspension 5.68 x 10^6^ particles/mL). A 5*μ*L sample was taken from the working solution to confirm the desired concentration by direct particle counting. Images acquired on a Ziess Axio Imager M2 were thresholded in ImageJ and the Particle Analyze function was used to count the number of microspheres present in the 5*μ*L sample. The solution was adjusted accordingly to a final concentration of approximately 4000 microspheres/100*μ*L. During surgery, prior to injection, the microsphere solution was vortexed for approximately 30 seconds to ensure microspheres were in suspension.

##### Surgical protocol

In preparation for the surgery, mice were placed in an isofluorane induction chamber (3.5% Iso in O_2_; 1L/min) and then transferred to a nose cone (2% Iso) and prepared for surgery by shaving the upper chest and neck, sterilizing the area and applying eye lubricant (Optixcare). Meloxicam was injected subcutaneously at 5mg/kg before transferring the mouse to a nose cone in the prone position on a heating pad set to maintain body temperature at 37°C.

A surgical incision at the neck (approximately 1.5cm) was created to gain access to the common carotid artery (CCA). Under a dissecting microscope, the internal carotid artery was carefully isolated from the Vagus nerve and looped loosely with a silk suture. A microvascular clip was applied to the external carotid artery to force the injected volume to travel into the internal carotid artery. To prepare for the injection, the free ends of the silk suture around the CCA were gently pulled to generate tautness in the vessel and a 31G needle was used to inject 100*μ*L of the microsphere solution (4000 microspheres/100*μ*L) over approximately 30 seconds. The microspheres stochastically distribute throughout the hemisphere and become lodged in penetrating arterioles. Once the needle is withdrawn from the CCA, surgifoam was applied to the injection site to control bleeding. The clamp from the external carotid artery was removed and the incision was sutured. Following the application of bupivacaine (2% transdermal, 0.1mL per animal) the mice were allowed to recover in a 30°C incubator prior to returning to their home cage.

#### Euthanization and vessel labelling

Mice were anesthetized with an intraperitoneal injection of sodium pentobarbital (65mg/mL, 10mL/kg). After confirming lack of responsiveness to a toe pinch, an incision was made to open the thoracic cavity. For visualization of the cerebral vasculature, 100*μ*L of Evans Blue (EB) (Sigma E2129, MW: 960.8g/mol, 2% solution in saline) was injected into the left ventricle of the heart with a 31G needle. The EB dye was allowed to circulate for a minimum of 5 minutes prior to extracting the brain. Brains were not perfused, in order to retain both microspheres and vascular labeling. Instead, tissue was post-fixed in 4% paraformaldehyde for 24 hours, and stored at 4°C.

#### Histological processing

Brains were sectioned at 100*μ*m thickness on a vibratome and sections were placed on 1% gelatin-coated slides. Slides were coverslipped with Fluoromount-G prior to imaging on a Ziess Axio Imager M2. For angiophagy quantification, individual microspheres (GFP; 510nm emission) and vessels (Evans Blue; 610nm emission) were imaged at 20x magnification. For brain-wide quantification and localization of microspheres, sections were tile-scanned at 2.5x (GFP; 510nm emission) and stitched using Zen software.

#### Microsphere localization and quantification

The previously defined semi-automated QUINT workflow was used for quantification and spatial analysis of microspheres on a brain-wide scale. The image analysis pipeline consists of three open-source software packages: Ilastik, QuickNII and Nutil. Raw image files acquired from the Ziess Axio Imager M2 at 10x were pre-processed with IrfanView (serial order and batch rename), Nutil (resize), and ImageJ (rotate). The Pixel Classification tool in Ilastik is used to generate segmentation files to differentiate the microspheres from the background. 10% of total acquired sections (approximately 7 images) were used to train the Pixel Classifiers in Ilastik. Annotations were applied to mark the microspheres against the background. Additional annotations were added to improve accuracy of predicted pixel classification, which is made based on pixel size, intensity and colour. The images were manually assessed and annotated until 100% accuracy was reached across all training images. The remaining serial sections were imported and the trained classifier generated segmentation maps for the whole series using the batch application feature. Batch processing in ImageJ was used to visualize segmentation files in colour with the glasbey lookup table. In QuickNII (RRID: SCR_016854), the acquired serial sections of the experimental brain are superimposed with the atlas (Allen Mouse Brain 2017) to allow for manual alignment, taking into account the size, position, orientation and sectioning angle. Predetermined landmarks are typically aligned first (development of corpus callosum, anterior commissures, and anterior hippocampus), followed by the remaining sections. The manually-adjusted serial alignments of the experimental and reference atlas sections is exported from QuickNII with an XML file describing a set of vectors regarding image position in relation to the atlas.

Microsphere quantification and localization was performed with the quantifier feature in Nutil. The Ilastik segmentation files, QuickNII alignment files, and the QuickNII XML file were imported into Nutil. Nutil then generated a list of objects (microspheres) with label IDs that correspond to the reference atlas and the object’s spatial coordinates. The final output from Nutil stated the number of microspheres in each brain region (cortex, fibretracts, hippocampus, olfactory, midbrain/hindbrain/medulla, thalamus, and ventricular system) as well as the size of the region. Microsphere distribution was normalized to region area by dividing the number of microspheres in a given region by the size of the region as determined by Nutil.

#### Angiophagy quantification

The timecourse of angiophagy was determined based on the location of microspheres in relation to the nearest visible vessel (labelled with Evans Blue) and was categorized into one of three classifications: obstructing, going out, or extravasated. Up to 20 microspheres (5 minimum) were quantified per brain region (cortex, striatum, hippocampus, thalamus and hypothalamus). Brains with fewer than 10 microspheres eligible for quantification were excluded from analysis.

#### Statistical analysis

All values were expressed as mean ± standard error of mean (SEM). All statistical analyses were performed in GraphPad Prism 10 (San Diego, CA, USA). One-way analysis of variance (ANOVA) followed by Tukey’s multiple comparison was used to analyze microsphere distribution across cohorts. Two-way ANOVA with Tukey’s multiple comparison was used to analyze microsphere localization across regions/cohorts and quantification of angiophagy. All p-values < 0.05 were considered significant. To analyze the correlation plots, Pearson’s bivariate correlation was used.

## Acknowledgements

This work was supported by a Canadian Institute for Health Research Project Grant, a Heart and Stroke Foundation of Canada Grant in Aid and a Canadian Foundation for Innovation grant to GS. FH was supported by a CGS-M scholarship, and the authors wish to tank Dr. Jessika Royea and Barman Loghmani for their assistance at various phases of this project.

## Supplemental figures

**Supplemental Figure 1.**
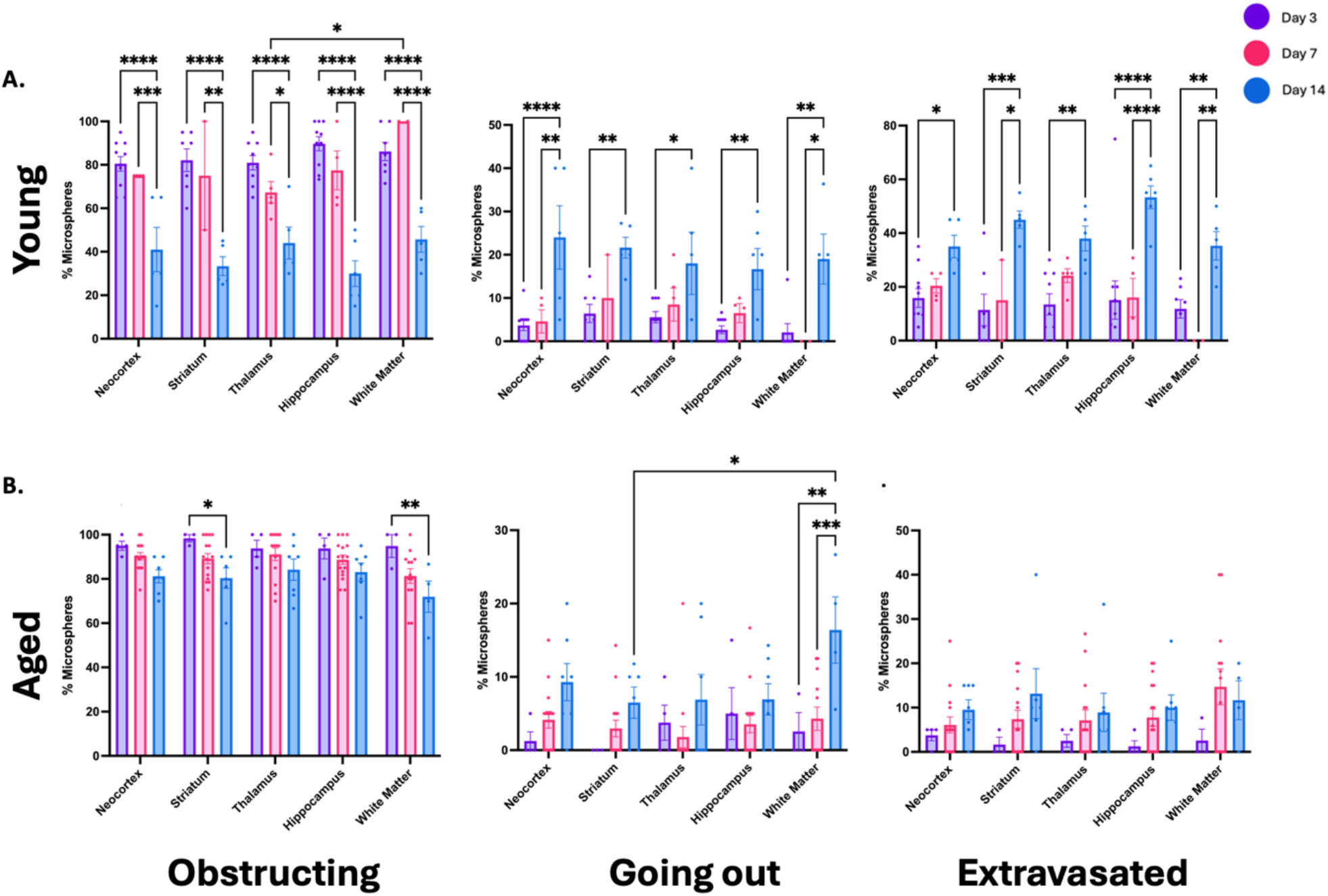
Quantification of angiophagy per brain region in young and aged non-diseased mice. Microspheres were quantified as obstructing the vessel, going out of the vessel, and extravasated from the vessel in each of five brain regions: neocortex, striatum, thalamus, hippocampus, and white matter at day 3, 7, and 14 for young (A) and aged (B) non-diseased mice. Data expressed as mean ± SEM. Two-way ANOVA with Tukey’s multiple comparison. ^****^p<0.0001, ^***^p<0.001, ^**^p<0.01, ^*^p<0.05.

**Supplemental Figure 2.**
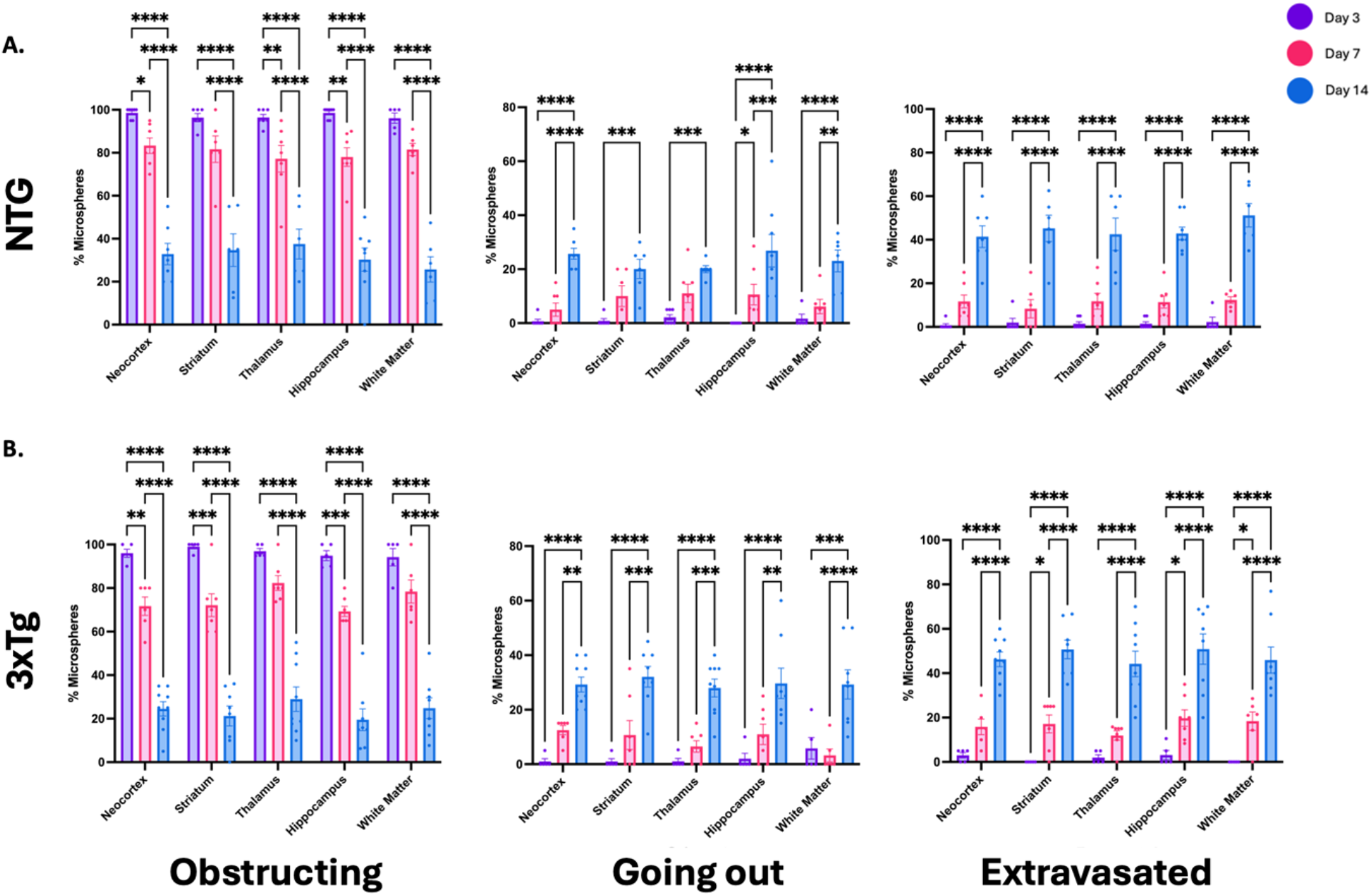
Quantification of angiophagy per brain region in young 3xTg and NTG mice. Microspheres were quantified as obstructing the vessel, going out of the vessel, and extravasated from the vessel in each of five brain regions: neocortex, striatum, thalamus, hippocampus, and white matter at day 3, 7, and 14 for NTG (A) and 3xTg (B) young mice. Data expressed as mean ± SEM. Two-way ANOVA with Tukey’s multiple comparison. ^****^p<0.0001, ^***^p<0.001, ^**^p<0.01, ^*^p<0.05.

**Supplemental Figure 3.**
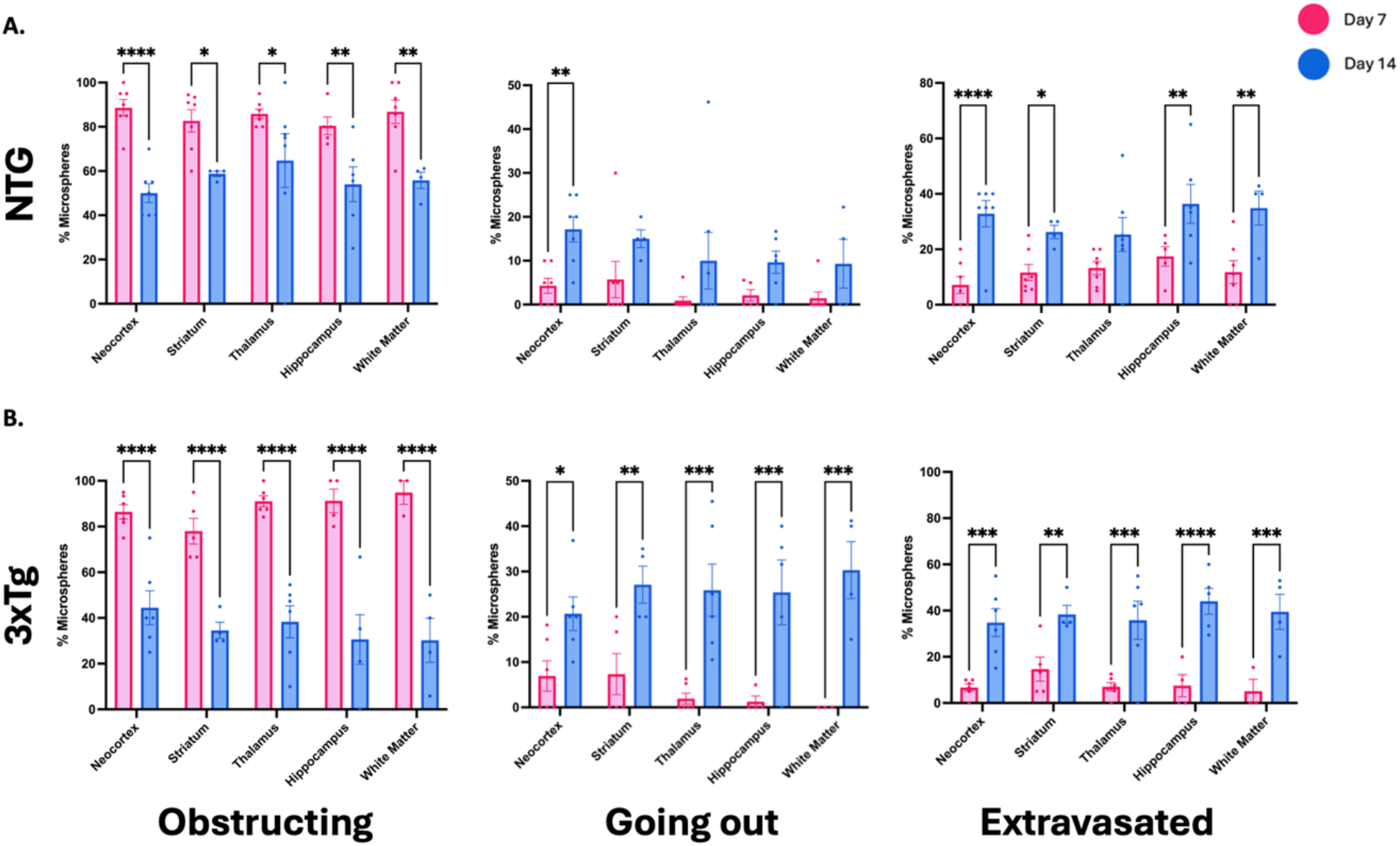
Quantification of angiophagy per brain region in aged 3xTg and NTG mice. Microspheres were quantified as obstructing the vessel, going out of the vessel, and extravasated from the vessel in each of five brain regions: neocortex, striatum, thalamus, hippocampus, and white matter at day 3, 7, and 14 for NTG (A) and 3xTg (B) aged mice. Data expressed as mean ± SEM. Two-way ANOVA with Tukey’s multiple comparison. ^****^p<0.0001, ^***^p<0.001, ^**^p<0.01, ^*^p<0.05.

